# Histone 3 lysine 9 dimethylation by the G9a-GLP heterodimer requires intranucleosomal product reading

**DOI:** 10.64898/2026.01.21.700667

**Authors:** Farzad Yousefi, Eric A Simental, Yongming Du, Sam Whedon, Michael J Trnka, Daniel Darling, Sohpia Jia, Barbara Panning, Philip A Cole, Mario Halic, Bassem Al-Sady

## Abstract

Repressive histone methyltransferases carry a catalytic (“write”) domain and a separate domain specialized for recognizing (“reading”) the reaction product. This read-write configuration acts as a positive feedback mechanism for epigenetic maintenance and the growth of repressive chromatin domains. Feedback exhibits as catalytic stimulation and is understood to act towards a proximal (*trans*) nucleosome. Whether this stimulation affects a specific methylation transition and whether it is restricted to trans-stimulation remains opaque. Here, we dissect the positive feedback in the heterodimeric histone 3 lysine 9 (H3K9) mono- and dimethlyase G9a-GLP, which carries two catalytic SET and two product-reading Ankyrin repeat (ANK) domains. We find that reading by both ANK domains is required for H3K9 di-, but not monomethylation on nucleosomes and for tight binding to them. As this read-writing occurs on dilute mononucleosomes, we propose that intranucleosomal feedback occurs for G9a-GLP. Swapping the ANK domains results in loss of dimethylation while maintaining nucleosome binding, indicating catalytic coupling of nucleosome methylation intermediates to reading. Crosslinking mass spectrometry reveals specific G9a surfaces that contact nucleosomal methylation intermediates. Structural approaches reveal how these surfaces position the G9a ANK domain on the methylation-intermediate nucleosome and stabilize G9a-GLP on chromatin during the reaction.

## INTRODUCTION

Repressive histone methyltransferases silence regions of the genome coding for lineage-inappropriate information as well as foreign genetic elements (1). Typically, these methyltransferases are initially recruited to a genomic locus via DNA-bound factors or RNA intermediates (2–4). Following this initial recruitment, domains carrying the repressive histone mark, for example, histone 3 (H3) lysine (K) 9 or 27 di (me2)- or trimethylation (me3) expand in a manner independent of the original DNA signal. This process is called heterochromatin spreading and is thought to require 3D contacts between product- and substrate nucleosomes mediated by the histone methyl-transferase (5, 6). Critically, it relies on the notion that all major repressive histone methyl transferases from Suppressor of variegation 3-9 (Suv39) type H3K9 trimethylases in *S. pombe* (7, 8) and animals (9, 10) to PRC2 H3K27 trimethylases (11) engage in product-positive feedback. The reaction product directs the enzyme further to substrate nucleosomes proximal in 3-dimensional space; thus, positive feedback is thought to operate in a trans-nucleosomal fashion. This is mediated by a separate “reader” domain in these enzymes beyond the catalytic site that is specialized for binding the reaction product. While the “reader” domains are members of different protein fold families, they share in common a methyl-lysine recognition mechanism based on an aromatic cage that engages in cation-π interaction (12). Whether product positive feedback in heterochromatin formation is implemented trans-versus intranucleosomally is not fully clear; however, trans-nucleosome engagement has been visualized for PRC2 (13).

Of the repressive histone methyltransferases, the H3K9 mono- and dimethylases G9a and GLP are unusual cases. These animal-specific paralogous enzymes are obligate dimers (14) that can homo (G9a-G9a; GLP-GLP)- or heterodimerize (G9a-GLP) and are essential for embryonic development in the mouse (14, 15). H3K9me2 domains produced by G9a/GLP assemble into lamin-associated domains at the nuclear periphery (16) and may serve as bookmarks through the nuclear envelope breakdown (17). Both enzymes contain a “reader” domain; in this case, the aromatic cage is embedded in an ankyrin repeat domain (18) and a catalytic SET domain common to all histone methyltransferases (19). Both enzymes carry an unstructured N terminus that serves as a recruitment platform to direct them to interaction partners, such as zinc finger transcription factors (20) or HP1 proteins. How the G9a and GLP ANK domains contribute to the formation of H3K9me2 domains is not fully clear, though it has been shown that G9a and GLP homodimers can be stimulated to methylate substrate nucleosomes in chromatin arrays that also carry product nucleosomes (21). This work also indicated that heterochromatin domain formation *in vivo* may not occur in a linear spreading fashion. However, homodimers are not the dominant form of the enzyme, especially in early embryonic tissues, where the heterodimer dominates (14). We have shown that there are significant biochemical differences between homo- and heterodimers in their ability to read product and engage in nucleosomal methylation, with the heterodimer significantly faster on chromatin than either homodimer (22). It is likely that homodimers have more specialized roles in the regulation of genes in differentiated cells (23) or in unique biological contexts such as oogenesis (24). Thus, how the heterodimer uses product reading in nucleosomal methylation remains unresolved.

Precisely how H3K9 methylases engage the nucleosome in the process of methylation and spreading remains unclear. The biochemical mechanisms that the “reader” domain uses in positive feedback have been described only in the context of the chromodomain. For Suv39/Clr4, the chromodomain engages on a H3K9me3 product nucleosome, which helps position the SET domain active site towards a substrate nucleosome linked *in cis*, elevating the catalytic rate (8) and mitigating nonproductive binding caused by tight nonspecific Suv39/Clr4 DNA association (25). For human Suv39h1, the chromodomain induces tighter binding to the H3K9me3 product nucleosome following initial recognition, decreasing the substrate K_M_ (9). The impact of product recognition on SETDB1 catalysis is less well understood (26, 27). While the structural basis for PRC2 stimulation by H3K27 methylation is well-evidenced (13), no similar structural insights into stimulation of H3K9 methyltransferases by H3K9 methylation have been forthcoming. How does product recognition change nucleosome binding and catalysis? Bulk binding is unaffected for Suv39/Clr4 (8); however, it is not clear if the enzyme changes its contacts with the nucleosomes following chromatin engagement. Is product binding coupled to methylation in some manner, for example, via allosteric changes? And if so, how are different methylation transitions affected? In several cases, product recognition elevates all catalytic transition rates from the unmethylated state (8, 11). Of note, higher accumulation of the terminal methylation state by feedback stimulation can be achieved both by elevating all transition rates or the last transition step specifically.

Here, we examine the role of product binding for the G9a-GLP heterodimer. We find that H3K9me1/2 recognition via the ANK domains is required specifically for the transition of H3K9me1 to me2 on the nucleosome. Since this occurs on mononucleosomes, this mechanism can operate intranucleosomally. G9a-GLP makes new contacts via reaction intermediate nucleosomes that bear a product tail, and these contacts appear to coordinate dimethylation. Finally, we provide insight into how the ANK domain engages a reaction intermediate nucleosome, showing that it binds across the nucleosome face, integrating contacts with the H4 and H2A N-terminal tails.

## RESULTS

### ANK domain aromatic cages are required for the H3K9 dimethyl transition on chromatin

We have previously shown (22) differences in H3K9 methylation on chromatin substrates between the three dimer forms of G9a/GLP. The G9a-G9a and GLP-GLP homodimers were slower and very slow, respectively, compared to the G9a-GLP heterodimer in methylating nucleosomes, while all three dimers had nearly indistinguishable kinetics on H3 peptide substrates. Specifically, the G9a-G9a homodimer appeared to have a similar ability to G9a-GLP to install H3K9 monomethylation (H3K9me1) but was noticeably slower in converting H3K9me1 to the dimethylated state (H3K9me2, **Figure 1A**). In addition to this kinetic defect, G9a-G9a homodimers were impaired in their ability to undergo “product recognition” i.e., bind to H3K9me1 or me2 peptides, while G9a-GLP can robustly bind both methylation states. Thus, the ability in G9a/GLP dimers to produce H3K9me2 correlates with the ability of the dimer to undergo product recognition. Here we test whether there is a mechanistic link between the two. Before analyzing this in nucleosomes, we performed Michaelis-Menten analysis of G9a-GLP on H3K9me0 and H3K9me1 peptides to understand if there is an intrinsic preference at the active sites. This analysis revealed only a minor difference in the Michaelis constant between H3K9me0 and H3K9me1 peptides, i.e., the K_M_ for H3K9me1 is moderately (50%) reduced (SFigure 1). Notably, a similar minor K_M_ increase by SET domain-only constructs of G9a (∼10%) or GLP (∼50%) for monomethylation was previously reported under saturating S-adenosyl methionine (SAM) concentrations (28).

**Figure 1:**
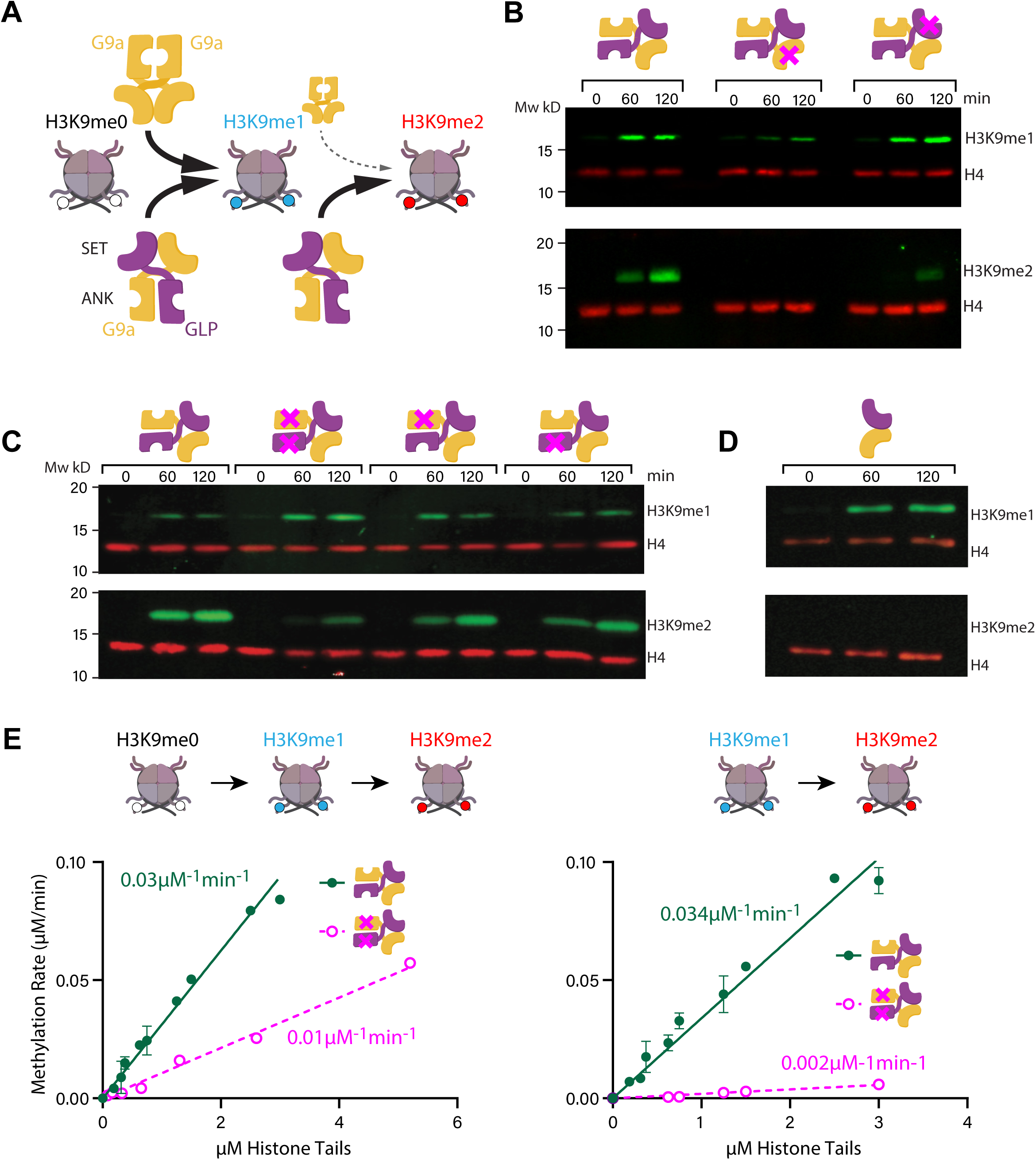
ANK domain aromatic cages are required for the H3K9 dimethyl transition on chromatin. **A.** Diagram of H3K9 mono- and dimethylation preferences on nucleosomes by G9a-G9a homodimer and G9a-GLP heterodimer, as shown in Sanchez et al. G9a-G9a is depressed in its ability to convert H3K9me1 to H3K9me2. **B.** Western blot of H3K9me1 and me2 production over time from H3K9me0 nucleosomes with wildtype G9a-GLP, and SET domain catalytic mutants G9a^SETm^-GLP and G9a-GLP^SETm^. **C.** As above, but for wildtype and ANK domain aromatic cage mutants G9a^ANKm^-GLP, G9a-GLP^ANKm^, and G9a^ANKm^-GLP^ANKm^. Note, the H3K9me1 antibody evidences some background in the t=0 timepoint. **D.** As above but for wildtype and deletion of G9a and GLP ANK domains **E.** Measurement of turnover rates on H3K9me0 (left) or H3K9me1 (right) nucleosomes under kcat/KM conditions (G9a-GLP concentration: 500 nM) with wildtype G9a-GLP or G9a^ANKm^-GLP^ANKm^. In this regime, the slope is roughly equivalent to the specificity constant.

Next, we established the role of either G9a or GLP SET domain in producing nucleosomal H3K9me1 or me2. To track the accumulation of methylation states on mononucleosomes, we take advantage of quantitative Western blots that allow the detection, on the material from the same reaction, of H3K9me1, H3K9me2, and H4 as a loading control, at t=0, 1, and 2 hours post initiation of methylation by the addition of S-adenosyl methionine.

We produced G9a-GLP dimers where either the G9a or the GLP SET domain is mutated at the active site and largely catalytically inactive (22, 29, 30), which allows us to assess the contribution of either SET domain to nucleosomes H3K9 mono- or dimethylation. Mutation of the G9 SET domain leads to a dramatic reduction in H3K9me1 accumulation and no H3K9me2 accumulation (**Figure 1B**). In contrast, the GLP SET mutant still allows wild-type levels of H3K9me1 production but is impaired in H3K9me2 accumulation, though not as strongly as the G9a SET mutant. We interpret this to indicate that both SET domains contribute to nucleosomal H3K9 methylation and that either 1. G9a is the dominant methylase, producing most H3K9me1, and with GLP making a meaningful but secondary contribution, or that 2. G9a is the primary H3K9 monomethylase, with GLP and G9a contributing equally to H3K9 dimethylation. The data cannot distinguish between these possibilities. Since H3K9 monomethylation appears to be fully unaffected in GLP SET mutants, we favor the second possibility.

Next, we examined how H3K9me product reading contributes to H3K9 methylation on nucleosomes. To do so, we mutated the aromatic cages of ankyrin repeat domains (ANK) that are required for H3K9me1 and H3K9me2 recognition (18, 22). We introduced ANK aromatic cage mutants (ANK^m^) into either G9a, GLP, or both in the G9a-GLP heterodimer. G9a ANK^m^ and GLP ANK^m^ in the heterodimer had a similar effect: H3K9 methylation is not discernibly affected, while the rate of H3K9 dimethylation is reduced (**Figure 1C**). However, when we mutate both ANK aromatic cages in the heterodimer, the resulting G9aANK^m^-GLPANK^m^ enzyme shows a significant increase in H3K9me1 and a strong reduction in the H3K9me2 accumulation. This result indicates that H3K9me1/2 reading is required for H3K9 dimethylation, and that both ANK domains contribute (**Figure 1C**). As there is residual H3K9me2 accumulation, we sought to determine the extent to which the ANK contributes to H3K9 dimethylation. As the G9a/GLP dimers are stabilized through the SET domain, deletion of the ANK leaves the major dimer interface intact. We deleted the entire ANK domain and observed that, while H3K9 monomethylation remains intact, H3K9 dimethylation is now completely abolished (**Figure 1D**). To more robustly quantify this effect, we performed methylation experiments with wildtype or G9aANK^m^-GLPANK^m^ starting with either H3K9me0 or H3K9me1 nucleosomes and measured initial rate methylation kinetics (**Figure 1E**). We note a ∼60% reduction in methylation when examining all S-adenosyl homocysteine release starting with H3K9me0 nucleosomes, but a more dramatic ∼95% reduction in the methylation rate when starting with the H3K9me1 nucleosome substrate. This indicates that the majority of the catalytic defect indeed occurs at the H3K9me1 to H3K9me2 transition and explains why, under western blot conditions, H3K9me1 accumulates to a higher degree in G9aANK^m^-GLPANK^m^ than in the wildtype. Importantly, neither the ANK deletion (ΔANK, SFigure 2B) nor G9aANK^m^-GLPANK^m^ (SFigure 2E) show any defect in mono-or dimethylating H3K9me0 H3 tail peptides when resolved by mass spectrometry, indicating no general impact on the SET domain in these mutants.

These experiments reveal that H3K9 dimethylation on nucleosomes by G9a-GLP, but not initial H3K9 monomethylation, requires the ANK domain, and recognition of H3K9me1/2 via the aromatic cages, specifically. This is in contrast to other repressive histone methyltransferases where product recognition allosterically, or otherwise (8, 9, 11), stimulates enzyme catalysis broadly.

### G9a/GLP ANK arrangement and aromatic cages are required for binding H3K9me1 and H3K9me2 on nucleosomes

We next wanted to assess if this catalytic enhancement is accompanied by tighter binding of the heterodimer to nucleosomes bearing methylation marks. The presence of methyl-reading domains in histone methyl transferases does not necessarily predict a change in bulk dissociation constant (K_d_), as is the case for Clr4 (8). This is because these enzymes often contain several DNA or histone octamer binding surfaces that are not methylation-specific. We assessed G9a-GLP: nucleosome interaction using electrophoretic mobility gel shift assays (EMSA), where the complex is stabilized under mild crosslinking conditions. First, we examined wildtype G9a-GLP on H3K9me0 or H3K9me2 nucleosomes. Nonetheless, by tracking the signal of the forming complex and the depletion of free nucleosomes, we can discern that G9a-GLP binds H3K9me0, albeit weakly. Tracking the disappearance of nucleosome shows clear binding > 2.5µM, indicating a K_1/2_ of 3.67 µM, similar to the K_M_ we had previously reported (**Figure 2A**)(22). Binding to H3K9me2 nucleosomes results in obvious complex formation at lower G9a-GLP concentrations (K_1/2_ = 0.56 µM), and about ∼6.5X preference for the H3K9me2 nucleosomes (**Figure 2A**). In addition, we clearly identify two G9a-GLP nucleosome complex bands, with the faster-migrating complex (CI) forming at lower G9a-GLP concentrations, and the slower-migrating complex (CII) appearing at higher concentrations. These two nucleosome-bound complexes were also detected with H3K9me1 nucleosomes, which appear to bind with similar affinity to the G9a-GLP heterodimer (**Figure 2B**).

**Figure 2:**
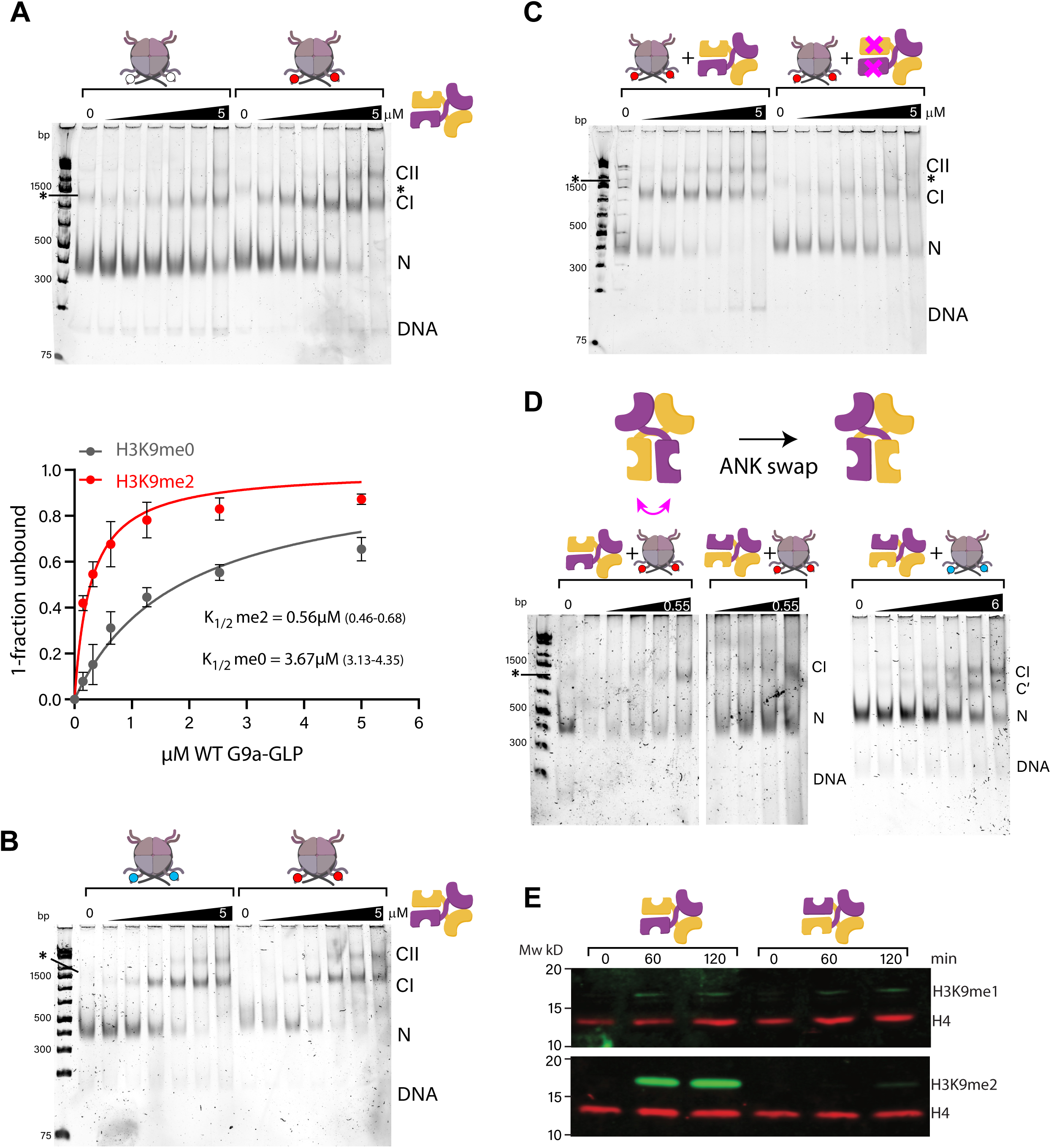
G9/GLP ANK arrangement and aromatic cages is required for binding H3K9me1 and H3K9me2 on nucleosomes. **A.** Electrophoretic mobility shift assay (EMSA) with H3K9me0 and me2 nucleosomes and G9a-GLP. Nucleosomes (15 nM) were incubated with G9a–GLP at concentrations (0, 0.15, 0.32, 0.63, 1.25, 2.5, and 5 µM) and subsequently crosslinked with 0.1% glutaraldehyde. The fitted K_1/2_ curves are shown below, indicating a specificity of 6.5X. A slight nonspecific band overlaps the G9a-GLP:nucleosome complex (asterisk). **B.** EMSA with H3K9me1 and me2 nucleosomes and G9a-GLP as in A. (including concentration regime). **C.** As above, but for H3K9me2 nucleosomes with wildtype G9a-GLPor G9a^ANKm^-GLP^ANKm^ as in A. (including concentration regime) **D.** TOP: Swap of the ANK domains of G9a and GLP in the heterodimer. BOTTOM LEFT: EMSA with me2 nucleosomes and wildtype G9a-GLP or G9a-GLP^SWAP^. G9a-GLP concentrations were 0, 0.067, 0.135, 0.27, and 0.55 µM. BOTTOM RIGHT: EMSA with me2 nucleosomes and G9a-GLP^SWAP^. G9a-GLP^SWAP^ concentrations for me1 nucleosomes were 0, 0.09, 0.19, 0.38, 0.75, 1.5, 3, and 6 µM. **E.** Western blot of H3K9me1 and me2 production over time from H3K9me0 nucleosomes with wildtype G9a-GLP or G9a-GLP^SWAP^ as in Figure 1. We note that in this experiment, the primary antibody detected a lower level of H3K9me1.

We suspected that this tighter, H3K9me1/2-dependent nucleosome binding is likely mediated by the ANK aromatic cages. To test this directly, we compared G9a-GLP:H3K9me2 nucleosome complex formation in wildtype versus G9aANK^m^-GLPANK^m^. We find indeed that the G9aANK^m^-GLPANK^m^ now behaves much more similarly to wildtype on H3K9me0 nucleosomes, i.e., unable to recognize H3K9 methylation (**Figure 2C**).

So far, we have shown that G9a and GLP ANK aromatic cages are required for H3K9 dimethylation and tight binding to H3K9 me1/2 nucleosomes. However, the inability of G9a-G9a homodimers to produce H3K9me2, or for G9a-G9a to recognize H3K9me1/2, prompted us to examine whether the native arrangement of ANK and SET domains, not just their presence, in the heterodimer is critical. To probe this, we generated G9a-GLP heterodimer with a different ANK-SET arrangement than wildtype without changing a single amino acid. Specifically, we “swapped” the ANK-core region of GLP with that of G9a and vice versa (see methods), fusing each to the reciprocal SET-containing region (G9a-GLP ANK^SWAP^). This “swap” preserves the wildtype complement of ANK and SET domains but just changes their orientation. We first tested how this G9a-GLP ANK^SWAP^ engages with H3K9me1/2 nucleosomes. In our hands, it is difficult to consistently express G9a-GLP ANK^SWAP^ to high levels. We performed a side-by-side binding experiment with wildtype to about the K_1/2_ point (0.55μM) on H3K9me2 nucleosomes and found that both wildtype and G9a-GLP ANK^SWAP^ bind roughly similarly (**Figure 2D**, left). We also tested G9a-GLP ANK^SWAP^ on H3K9me1 nucleosomes and were able to prepare a more concentrated prep for this experiment. This experiment shows clearly that G9a-GLP ANK^SWAP^ engages H3K9me nucleosomes, though with somewhat lower apparent affinity than wildtype, and forms a characteristic secondary complex we observe on me1/me2 nucleosomes for wildtype (**Figure 2A-C**) at higher concentrations. Interestingly, this complex migrates at a different apparent size in G9a-GLP ANK^SWAP^ than for the wildtype.

We next tested in G9a-GLP ANK^SWAP^ has a catalytic defect similar to G9aANK^m^-GLPANK^m^ in nucleosomal H3K9 dimethylation. Indeed, G9a-GLP ANK^SWAP^ has an even more severe phenotype, with H3K9 dimethylation now completely abolished, whereas H3K9 monomethylation remains unaffected. As for G9aANK^m^-GLPANK^m^, G9a-GLP ANK^SWAP^ does not show any defects in producing H3K9me2 on H3 tail peptides (SFigure 2C). As with G9aANK^m^-GLPANK^m^, G9a-GLP ANK^SWAP^ does not show any defects in producing H3K9me2 on H3 tail peptides (SFigure 2C). These results indicate that the ANK domain aromatic cages and their specific orientation relative to the two SET domains are crucial for nucleosome H3K9 dimethylation.

### G9a-GLP makes unique contacts with a reaction intermediate, involving multiple contacts in the ANK domain

Since the above data suggest that the ANK makes critical contacts with the nucleosomes required for H3K9 dimethylation, we were interested in probing if such specific contacts arise during the reaction cycle. To examine how G9a-GLP may interact with an initial unmethylated substrate and a reaction intermediate, we produced corresponding substrate mimics for structural analysis. To mimic substrate H3 tails, we replaced the target lysine with a linear aliphatic side chain, which has been shown to “trap” the histone tail in the catalytic SET domain active site. Prior structural work has made use of K to M mutations for the study of PRC2 (31) and G9a (32), as well as for non-natural substitutes, such as ethylcysteine (EcX) (33) and norleucine (Nle) (34) (SFigure 3A). We systematically compared substrate mimics here and found that both EcX and Nle substitutions on the nucleosome bind G9a-GLP comparably, while a H3K9M nucleosome binds more weakly in our hands (SFigure 3B). To produce a reaction intermediate that mimics one tail in the H3K9me0/1-substrate state and the other in the H3K9me2 product state, we produced an asymmetric nucleosome containing one H3K9Nle tail (**Figure 3A**) and a second H3K9me2 tail.

**Figure 3.**
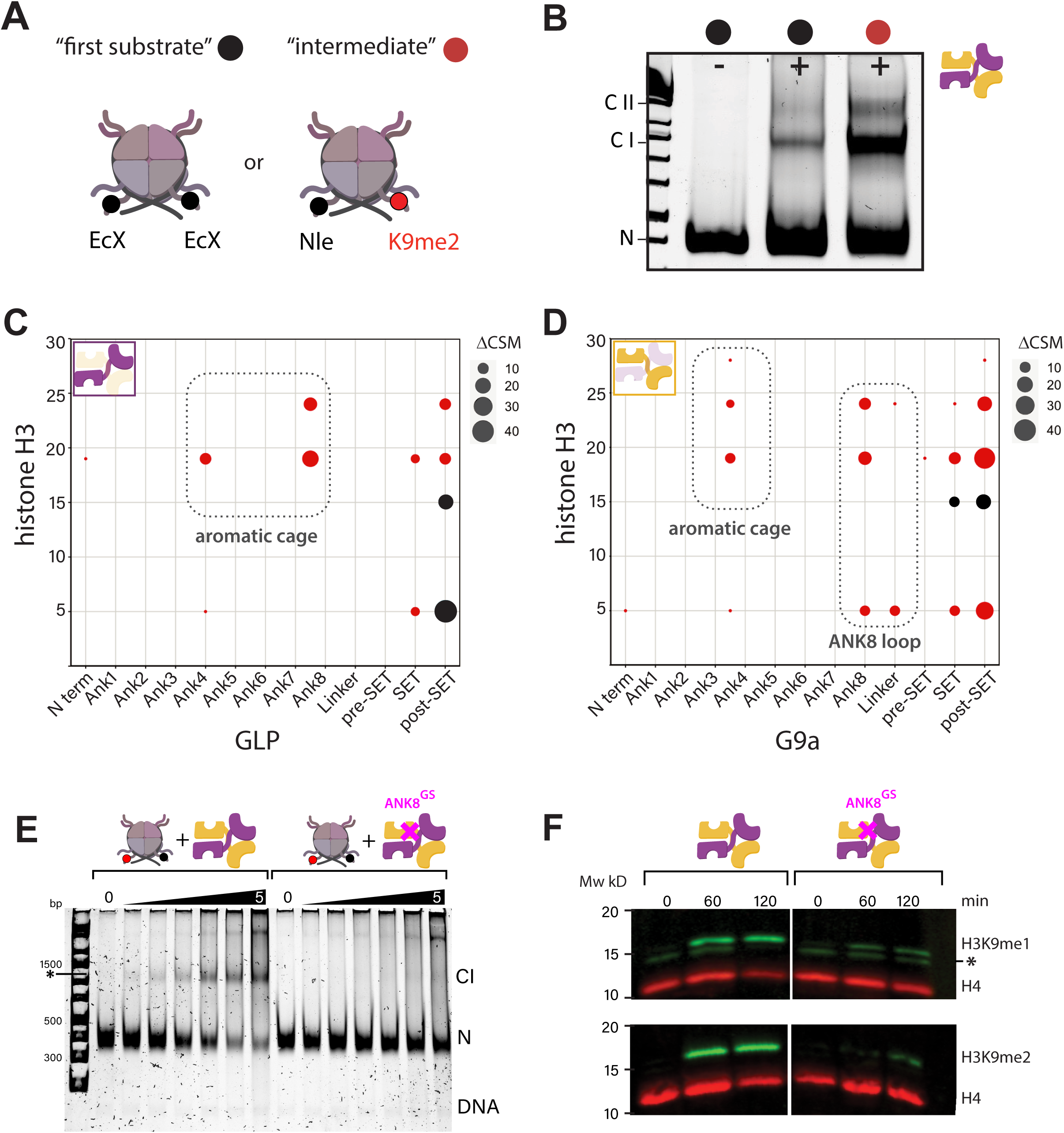
G9a-GLP makes unique contacts with a reaction intermediate, involving multiple contacts in the ANK domain. **A.** diagram of initial substrate and reaction intermediate mimics. The reaction intermediate mimic is asymmetric H3K9Nle/H3K9me2. **B.** complex formation for crosslinking mass-spectrometry (CLMS). Lane 1, H3K9Ecx mononucleosomes; lane 2, G9a–GLP bound to H3K9Ecx “initial substrate” mononucleosomes; lane 3, G9a–GLP bound to asymmetric H3K9me2/H3K9Nle “reaction intermediate” mononucleosomes. Samples were crosslinked with DSSO (800 µM) prior to electrophoresis (see Methods). **C.** CLMS overview crosslinked peptides from the initial substrate mimic (black) and reaction intermediate mimic (red) to GLP. Only crosslinks to H3 tail are shown. **D.** As in C., but for crosslinks to G9a. **E.** EMSA with reaction intermediate mimic nucleosomes and wildtype G9a-GLP or G9aANK8^GS^-GLP. **E.** Western blot of H3K9me1 and me2 production over time from H3K9me0 nucleosomes with wildtype G9a-GLP or G9aANK8^GS^-GLP as in Figure 1.

To pinpoint the interactions of our initial substrate and reaction intermediate mimics with G9a-GLP, we performed crosslinking Mass Spectrometry (CLMS) (see methods). Following crosslinking, EMSA of G9a-GLP-nucleosome complexes suggested stronger complex formation with the reaction asymmetric intermediate mimic as compared to the initial substrate mimic (**Figure 3B**), which is in line with the stronger association with H3K9me2 versus me0 nucleosomes we document above. CLMS detected unique contacts of different parts of G9a-GLP with either substrate (**Figure 3C**, D and SFigure 3C, D). Specifically, G9a and GLP contacted the initial substrate mimic only at the H3 tail via the SET or post-SET domain, indicating that the first contacts are dominated by a catalytic site engagement. In contrast, both G9a and GLP made expanded contacts with the tail. While both G9a and GLP similarly made contacts at the SET and post-SET domain associated with catalysis, both enzymes also contact the reaction intermediate mimic near their respective aromatic cage binding pockets, constituted by residues in ANK 4,5 (18) but proximal to residues in ANK 3, 6, and 7. In addition, G9a contacts the H3 tail in a region just downstream, namely ANK8 and the linker between the ANK and SET domains. Further, the N-termini of both G9a and GLP contact the H2A tail, while the ANK8 region of G9a also contacts the H4 tail (SFigure 3C, D). We examined more closely the sequence in the region where the ANK8:nucleosome crosslink occurs in G9a and, surprisingly, found a uniquely non-conserved region between GLP and G9a. The G9a sequence (887–894) is AWDLTPER, the GLP sequence (975–982) is PLQCASLN, with the target crosslinking lysine a few amino acids upstream (SFigure 4A). To test if this region, which makes unique H3 and H4 tail contacts to the reaction intermediate, has functional significance, we mutated it in G9a to a glycine-serine (GS) linker. This ANK8 GS mutant in G9a (G9aANK8^GS^-GLP) methylates H3 tail peptides similar to wildtype and SWAP and G9aANK^m^-GLPANK^m^ mutants, again indicating no change in intrinsic catalytic capacity (SFigure 2F). However, the situation on the chromatin substrate is different: Side-by-side with wildtype G9aANK8^GS^-GLP binds much more poorly to the reaction intermediate mimic (**Figure 3E**), as well as to H3K9me1 nucleosomes (SFigure 4B). Consequently, G9aANK8^GS^-GLP is compromised in methylation on the nucleosome: Both H3K9 monomethylation and dimethylation are impacted; however, dimethylation is more severely reduced (**Figure 3F**). This indicates that the ANK8 of G9a critically coordinates the binding of the reaction intermediate to facilitate further nucleosomal methylation, ultimately leading to the completion of symmetric H3K9 dimethylation as the final product.

### The ANK domain contacts the reaction intermediate across the nucleosome surface and additionally contacts the H4 and the H2A N-terminal tails

We next aimed to obtain structural insight into how G9a-GLP engages a reaction intermediate on its pathway to complete H3K9 dimethylation on the nucleosome. To do so, we performed cryo-electron microscopy on mildly glutaraldehyde-cross-linked complexes of G9a-GLP and the H3K9Nle/H3K9me2 reaction intermediate mimic. We purified the complexes from the binding reaction by size-exclusion chromatography (see SFigure 5A) and pooled the fractions that retained the highest proportion of bound nucleosome (fractions 3 and pooled fractions 4-6). Both fractions were combined and used for cryo-EM grid preparation. Initial classification revealed a very undefined density on the nucleosomal disc face (SFigure 5B). After extensive classification, we obtained two maps of a nucleosome with an additional density at an overall resolution of 7Å and 5.1Å, respectively. The additional density bound to nucleosome (**Figure 4A**, **C**; SFigure 5C, D) is significantly smaller than would be expected for the G9a-GLP ANK-SET dimer. Although the resolution of additional density is limited to 10Å-20Å and insufficient to resolve secondary structure, the available crystal structures of both G9a and GLP SET domains (19, 32) and ANK domains (18) allowed us to model either of these structures into the extra density. The extra density is strongly compatible with the ANK domain of either G9a or GLP (**Figure 4E**, G9a shown). Because their predicted structures are very similar, we cannot clearly distinguish which one is bound in the electron density map. However, we favor the ANK of G9a for the following reason: While both G9a and GLP ANK aromatic cages contact the H3 tail (**Figure 3**), only the G9a ANK8 shows cross-links to the H4 tail basic patch. Using two electron density maps (map 709, resolution 7Å; map 808, resolution 5.1Å, **Figure 4A,B**; SFigure 5C,D), we resolve two anchor points of the ANK on the nucleosome surface: The N terminus of the ANK makes contact with the H2A N-terminal tail (**Figure 4B**). As this tail is often found to reside in the DNA minor groove (35), it might swing out to engage the ANK. A separate map clearly shows electron density at the H4 tail (**Figure 4D**). A model produced from the maps using the G9a ANK domain (**Figure 4E**) shows that a key part of the contact to the H4 tail is the ANK8 unique region. This contact is validated by our CLMS data (SFigure 3C). The G9a unique sequence (SFigure 4A) is colored in light blue in Figure 4D, E, and G. We also observe interaction with the acidic patch of the nucleosome. Thus, in our model (**Figure 4E**), the ANK sites across the entire nucleosome disc face in a crescent moon shape. However, given the known flexibility and size of the H3 tail, we were unable to observe clear electron density for it. The ANK aromatic cage, positioned near the middle of the ANK domain, falls over the middle of the nucleosomal disc in our model. We do observe fuzzy electron density projecting from the exit of the H3 tail from the dyad toward the aromatic cage. The possibility that this represents an H3 tail looping back from the H3 tail exit site toward the ANK is supported by 1. CLMS data supporting H3 tail crosslinks to the vicinity of the ANK aromatic cage (crosslinks at H2B residues 100-117) and on the trajectory to the ANK aromatic cage in the reaction intermediate nucleosomes (crosslink at H2A C-terminus). These crosslinks are more numerous in the reaction intermediate complex than substrate mimic complex (SFigure 6B-D vs F-H), and some additionally are unique. In contrast, intranucleosomal crosslinks of the H3 core are more numerous in the substrate mimic (not shown). 2. Destabilization of the H3 αN helix and DNA unwrapping next to the H3 that is more proximal to the bound ANK, supporting further the notion that this tail is looping back toward the ANK. However, we stress that this data supports, but is not definitive proof of occupancy of H3K9me2 in the ANK aromatic cage in this structure.

**Figure 4:**
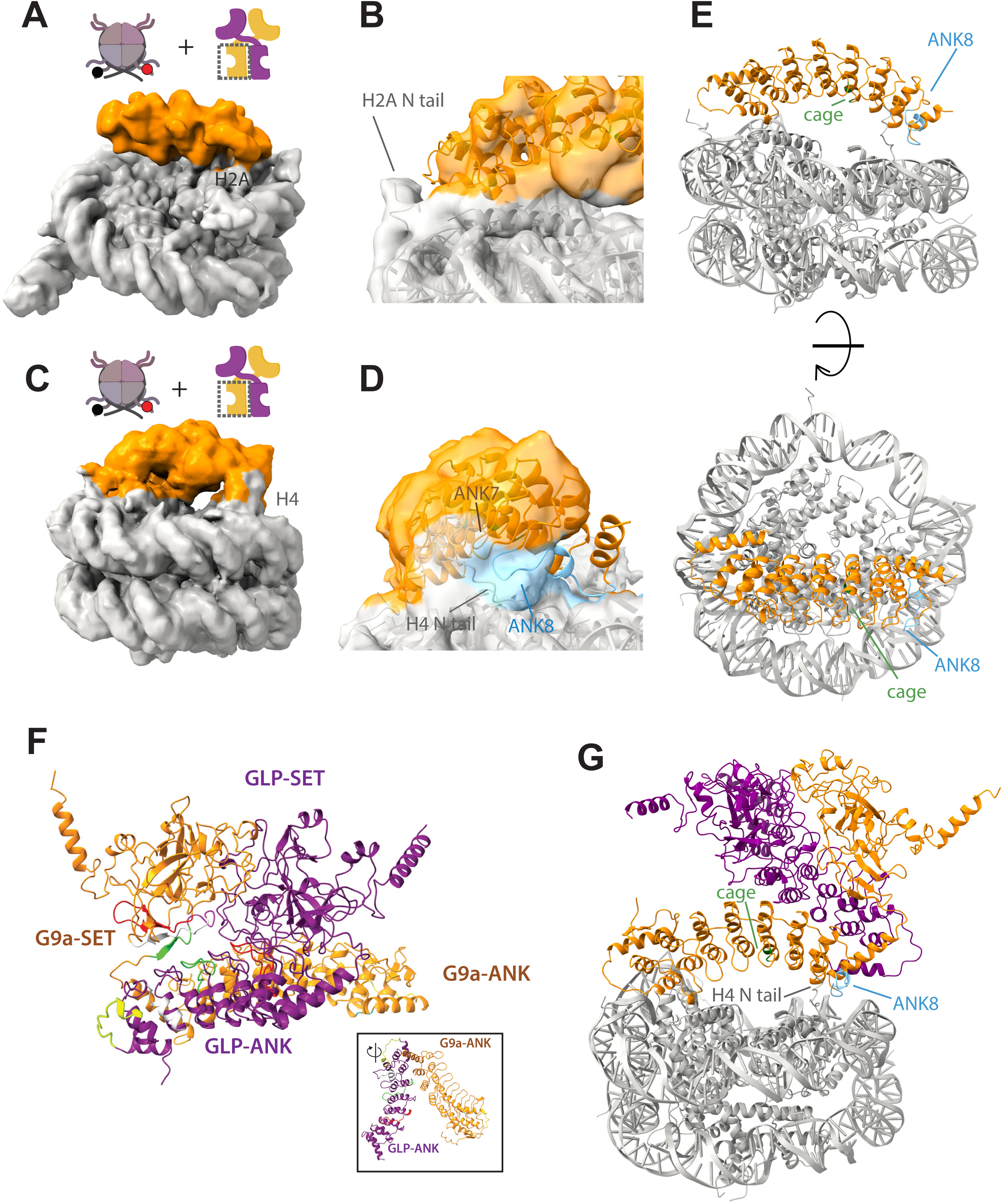
The ANK domain contacts the reaction intermediate across the nucleosome surface and additionally contacts the H4 and the H2A N-terminal tails. **A.** Side-view of cryo-EM density (map 709) of intermediate (H3K9Nle/H3K9me2) nucleosome bound by G9a ANK domain, nucleosome is colored in gray and G9a in orange. **B.** Detail of A. showing the H2A N-terminus:G9a ANK interaction. **C.** Side-view of cryo-EM density (map 808) of intermediate (H3K9Nle/H3K9me2) nucleosome bound by G9a ANK domain, colors as in A. **D.** Detail of B. showing the H4 N-terminus:G9a ANK7/8 interaction. The ANK8 unique sequence in G9a is colored in light blue. **E.** model combining all electron density maps of G9a and the ANK intermediate (H3K9Nle/H3K9me2) nucleosome. The ANK8 unique sequence in G9a is colored in light blue, and the ANK aromatic cage in green. **F.** AlphaLink2 model of the ANK-SET G9a-GLP heterodimer. Crosslinks are indicated with a unique color per crosslink. Inset: A rotated view of the G9a and GLP ANK domains. **G.** Structural model of G9a/GLP dimer bound intermediate (H3K9Nle/H3K9me2) nucleosome generated by X-MS and cryo-EM models, nucleosome is colored in gray, G9a colored in dark orange and GLP in purple.

Since the extra density covers the ANK, which we thus think represents the stably bound part of the G9a-GLP dimer, we were curious to model how the rest of the enzyme might sit in this orientation on the reaction intermediate mimic. To do so, we first produced a model of the G9a-GLP ANK SET dimer. Using the crosslinks of the enzyme in the CLMS experiment with the reaction intermediate nucleosome (S Figure 7A, B), we produced an Alpha-Link2 model. Alpha-Link2 directly incorporates experimental crosslinks into the open source UniFold protein structure modeling engine (36). The resultant model (**Figure 4F**) has generally high confidence and low prediction alignment error (SFigure 7C-E) and is superimposable with an independently derived AlphaFold3 model (SFigure 7F). In this AlphaLink2 model, the G9a-GLP SET domain heterodimer is similar to what has been previously described for either homodimer (19). The ANK domains splay apart from one another (inset), with the aromatic cages solvent accessible. We next docked the G9a-GLP AlphaLink2 model onto our nucleosome model of the G9a ANK. This allowed us to position the G9a-GLP ANK-SET dimer relative to the nucleosomes (**Figure 4G**). In this model, the SET domains of G9a and GLP are positioned above the ANK. Whether the H3 tail that is not engaged with the ANK reaches the SET domain active site in this orientation is unclear, but it may be that the writing step requires a rearrangement of the complex. In any case, we conclude that this model represents a “reading” orientation of G9a-GLP that we believe is a required intermediate step for the completion of nucleosomal dimethylation.

## DISCUSSION

Repressive H3K9 methylases can produce up to three methylation states: H3K9me1, me2, and me3. Of those, H3K9me1 is non-repressive (37, 38), H3K9me2 can be repressive in the right context (16, 39, 40), and H3K9me3 is generally repressive (41). G9a-GLP mostly produces non-repressive H3K9me1 and conditionally repressive H3K9me2. We show here that the transition to H3K9me2 is licensed by nucleosomal reading of an intermediate methylation state, which may be H3K9me1 or H3K9me2 on one of the two tails. This to our knowledge is a unique feature of G9a-GLP. The following three general principes derive form this work: (1) a specific methylation state can be boosted by stimulation of one transition, rather than general enhancement of all rates; (2) this stimulation can occur in an intranucleosomal manner, not just in a transnucleosomal fashion; and (3) Product reading by H3K9 methylases appears to occur on the nucleosomal disc, with the H3tail folding back into the aromatic cage.

### 1. Nucleosomal stimulation of H3K9me1 to me2 transition

Product stimulation is widespread in repressive histone methyl transferases, and is present in H3K9 methylases Clr4 (7, 8), Suv39h1 (9), and G9a/GLP (21), and to some degree for SETDB1 (27) and the H3K27 methylase PRC2 (11). In the case where individual transition rates have been measured directly, positive feedback stimulation elevates all rates, i.e., via overall increased activity at the active site. Similar stimulation via alternative mechanisms, such as ubiquitination, reveals broad activation via allosteric changes at the active site (42, 43). Since direct transition rates have not been measured in several cases, it is unclear whether product stimulation affects a specific rate. To our knowledge, no case has been documented in which only one specific transition is altered. Here, we document for G9a-GLP that nucleosomal product reading specifically enhances the H3K9me1 to me2 rate. This is peculiar, as on the H3 peptide, the difference between both transitions is relatively minor, with similar k_cat_ and a roughly doubled K_M_ for H3K9me1 versus H3K9me0 (SFigure 1). As none of the G9a-GLP mutants that impact nucleosomal dimethylation are affected in their ability to dimethylate H3 peptide, this stimulation of the me1-me2 transition is nucleosome-specific. What may underlie this nucleosome-specific effect? We posit two mechanisms: a. It may be the case that the H3 tail is differentially accessible in the H3K9me0 and me1state. The H3 tail associates with nucleosomal DNA (44). Unlike acetylation, which induces charge neutralization, H3K9 methylation does not induce large electrostatic changes, it has recently been shown that the H3K9 methylation state alters behaviors such as apparent nucleosome spacing and radius of gyration on chromatin arrays *in vitro* (45). Subtle changes may be enough to exaggerate the subtle K_M_ difference already apparent on the peptide on the nucleosome. Engagement with one H3K9me1 or me2 tail may then stabilize the enzyme on the nucleosome. b. Alternatively, it is possible that ANK engagement relieves an inhibition of dimethylation of the monomethyl H3K9 that is apparent on the nucleosome. The precise arrangement of the active site of H3-targeted SET domains determines the methylation state distribution via the F/Y switch (46). At the conserved position, G9a/GLP has the F residue, more compatible with di-and trimethylation. It may be the case that an adverse interaction with other parts of the H3 tail upstream from the target lysine or other interactions near the SET domain induces a conformation that disfavors a dimethylation-compatible active site conformation. ANK engagement may overcome this by either increasing residence time as above or transmitting an allosteric change that relieves this inhibition. Since the baseline catalytic preference is minor for H3^1-15^K9me0 versus K9me1peptides (SFigure 1) we favor a dimethylation inhibition model. Of note, a CLMS experiment we performed with H3K9me1 nucleosomes (not shown) showed several more interactions between G9a-GLP and the nucleosome than observed with either the initial substrate (K9Ecx) or reaction intermediate (K9Nle/K9me2) nucleosome.

### 2. Stimulation can occur in an intranucleosomal fashion

Product positive feedback in repressive histone methyltransferases is generally thought to promote transnucleosomal methylation. This has been directly shown for PRC2 (13) and is known to occur for Clr4 and Suv39h1 (8, 9). A study demonstrated that, for G9a homodimers and GLP homodimers, transnucleosomal stimulation may occur when nucleosomes are fully pre-modified with methyl lysine analogs Kc9me1 or Kc9me2 on chromatized plasmids (21). We did not observe transnucleosomal stimulation on dinucleosomes or linear arrays with homodimers (not shown), indicating that more complex 3D interactions present on circular templates may be required to induce this stimulation. However, the catalytic response of G9a-GLP to existing methylation has not been previously measured. The observation that product reading via the ANK is absolutely required to produce dimethylation in a dilute solution of mononucleosomes indicates that intranucleosomal stimulation of H3K9 methylation must occur. This, however, does not preclude the same mechanism applying across nucleosomes as well. In fact, the arrangement of ANK and SET domains captured in our combined model of cryo-EM density, crosslinking mass spectrometry data, and AlphaLink2 modeling indicates that the SET domains are distal to the substrate H3 tail (**Figure 4F**). It may be that this model does not capture the native domain arrangement or an allosteric change in the protein introduced by ANK engagement, which might trigger a large rotation. Intranucleosomal stimulation raises the question of whether the transition from mono- to dimethylation is gated locally at the nucleation site rather than distally via spreading in G9a-GLP. In fact, G9a and GLP are tightly associated with Zinc Finger transcription factors (ZnF) that recruit the enzymes (20, 47–49). It may be that product reading, possibly in addition to recruitment, together promote di- or even trimethylation, which then enables recruitment of silencing factors. The role of transnucleosomal stimulation by G9a-GLP warrants further investigation.

### 3. Nucleosomal product reading by a H3K9 methylase

Here, we present the first structure of an H3K9 methylase on a nucleosome. The distance of H3K9 from the nucleosome core and the flexibility of the H3 tail may explain our inability to resolve its interaction with either SET domain or ANK aromatic cage. This is in contrast to cases such as H3K27 methylase PRC2 (13, 50), H3K79 methylase Dot1 (34), or euchromatic methylases like COMPASS (51), where the target residue of H3 has been resolved along with a few adjacent residues. Instead, this work focuses on capturing the reading orientation of the G9a-GLP enzyme that is required for the production of the terminal H3K9me2 state on the nucleosome.

Our low-resolution electron density maps most likely represent the ANK domain of G9a engaging the nucleosome. Rather than seeing the ANK near the exit of the H3 tail, it resides in a crescent shape across the nucleosome surface. We believe that reading H3K9me1/2 is coupled to recognition of other nucleosomal elements: 1. the acidic patch, 2. the H2A N-terminal tail, and 3. the H4 tail near the basic patch (H4K16-K20). This provides an opportunity for both a more stable association and, possibly, regulation via post-translational modifications of H2A and H4. It is conceivable that H4K16 acetylation may repel the ANK domain, in a manner reminiscent of the SIR3 BAH domain (52), though this remains to be tested. Why do we not directly visualize the H3K9 residue in our density maps? This is likely because the H3 tail is long and disordered, only entering the nucleosome around residues P38-R40. This suggests that the H3 tail flips back across the dyad and follows a path across the nucleosome disc, along the dyad axis, into the aromatic cage of the ANK. CLMS data, fuzzy electron density of the H3 tail, destabilization of the αN of H3, and local DNA unwrapping support this view.

In our model of the G9a-GLP dimer on the nucleosome, the SET domains are not proximal to the nucleosome core. We present three possible interpretations for this observation: The SET domain, when anchored on the nucleosome disc by the ANK, may methylate the second H3 tail, given its length, in a manner compatible with the orientation we show. Alternatively, the model represents an initial contact that induces a conformational change in the enzyme, bringing the SET domain closer to the second H3 tail. This possibility is supported by unique cross-links in G9a-GLP with the reaction intermediate mimic, but not substrate mimic nucleosome (SFigure 7). These cross-links do not drive major changes in the Alpha-Link2 model, but this model does not use all available cross-links. Third, the conformation we capture may correspond to a read-write orientation relative to a neighboring nucleosome. Structural resolution on the substrate-mimic nucleosomes and polynucleosome substrates will allow distinction between these possibilities.

## MATERIALS AND METHODS

### Purification of G9a–GLP

The ANK–SET G9a–GLP complex was expressed and purified as described by Sanchez *et al*. Briefly, N-terminally His-tagged G9a and MBP-tagged GLP were coexpressed from a single bicistronic vector (QB3 Berkeley Macrolab) in *E. coli* Rosetta (DE3) cells. The complex was purified using sequential cobalt- and amylose-affinity chromatography, followed by TEV protease-mediated tag removal and size-exclusion chromatography on a Superdex 200 Increase 10/300 column. Purified protein was buffer-exchanged into storage buffer containing 100 mM Tris-HCl (pH 8.0), 100 mM KCl, 10% glycerol, 1 mM MgCl₂, 20 µM ZnSO₄, and 10 mM β-mercaptoethanol. ANK aromatic cage mutants and SET domain catalytic mutants were as in Sanchez *et al.* The G9a-GLP^SWAP^ was produced as follows: The ANK-core region of GLP (residues 107–376) was replaced with that of G9a (residues 50–322), and vice versa, with each fused to the reciprocal SET-containing region (G9a residues 323–613; GLP residues 377–667) to generate G9a-GLP^SWAP^ (ANK(GLP)–SET(G9a) and ANK(G9a)–SET(GLP)).In both constructs, the junction was placed at the conserved PIPCV motif. For G9aANK8^GS^-GLP, the ANK8 sequence in G9a AWDLTPER was replaced with GGGSGGG.

### Preparation of Mononucleosome Substrates

H2A, H2B, H3 C110A, and H4 histone proteins, as well as reconstitution for nucleosomes except asymmetric nucleosomes (see below), were prepared as described in Kennedy et al. (53). Asymmetric nucleosomes containing H3K9Nle/me2 were prepared as in (54). Briefly, synthetic H3.1 peptides (amino acids 1–34) incorporating either H3K9Nle or H3K9me2, were synthesized by Fmoc-based solid-phase peptide synthesis and purified via reverse-phase C18 HPLC. Recombinant histone octamers containing H2A, H2B, H4, and truncated H3(33–135) were co-expressed in *E. coli* and purified using Ni2+ affinity and size-exclusion chromatography. Widom 601 DNA templates (147 and 185 bp) were prepared by plasmid extraction, EcoRV digestion, and anion-exchange purification. Nucleosomes were reconstituted by combining histone octamers and DNA, followed by linear salt gradient dialysis (starting buffer: 2 M KCl, 10 mM Tris pH 7.5, 1 mM EDTA, 1 mM DTT; final buffer: 150 mM KCl, 10 mM Tris pH 7.5, 0.1 mM EDTA, 1 mM DTT; 36 hour linear gradient, followed by a final 3 hour dialysis against the final buffer). Sequential, site-specific incorporation of H3K9Nle and H3K9me2 was accomplished by sortase ligation using cW11 sortase and the corresponding synthetic peptide. Ligation of K9Nle was performed first (25 µM peptide, 1.65 µM nucleosome, 200 µM sortase, 50 mM Tris pH 7.5, 150 mM NaCl, 5 mM CaCl2, 37 °C, 4 hours), after which concentration of NaCl in the reaction was adjusted to 250 mM with 5 M NaCl, and nucleosomes containing one copy of H3(33–135) and one copy of H3K9Nle were purified by weak anion-exchange chromatography (TSKgel DEAE-5PW, 7.5 cm x 7.5 mm, 10 µM particle size, Tosoh; 147 bp gradient: 150 to 250 mM KCl in 10 mM Tris pH 7.9, 0.5 mM EDTA over 10 minutes, followed by 370 to 440 mM KCl over 21 minutes; 185 bp gradient: 150 to 250 mM KCl in 10 mM Tris pH 7.9, 0.5 mM EDTA over 10 minutes, followed by 430 to 480 mM KCl over 21 minutes) to remove unligated nucleosomes and truncated byproducts arising from cleavage of the T11-G12 bond uniquely observed with H3K9Nle peptides. The purified nucleosomes were dialyzed into the final dialysis buffer of the linear salt gradient, then subjected to a second round of the sortase ligation reaction using a higher concentration of H3K9me2 peptide (50 µM peptide), and no other alterations to reaction or purification conditions. Purified nucleosomes were dialyzed into 25 mM NaCl, 10 mM Tris, 1 mM DTT, 20% glycerol, concentrated to ∼5 µM and used immediately, or flash frozen and stored at −80 °C. Incorporation of histone modifications was verified by western blotting and LC-MS. Commercial mononucleosomes containing H3K9me1 (SKU 16-0325) and H3K9me2 (SKU 16-0324-20) were purchased from EpiCypher and used directly. H3K9Ecx histones were prepared as follows: Histone H3K9C was dissolved in isopycnic alkylation buffer containing 1.5 M HEPES (pH 8.0), 20 mM *DL*-methionine, and 20 mM dithiothreitol and incubated at 37 °C for 1 h to reduce cysteine residues. Alkylation was initiated by the addition of bromoethane (100 µL of a 1 M solution) and allowed to proceed for 4 h at room temperature. Excess alkylating reagent was subsequently quenched by the addition of dithiothreitol (10 µL of a 1 M solution) and incubated for >10 h at room temperature. The reaction was finally quenched with β-mercaptoethanol (50 µL, 14.2 M). Formation of H3K9Ecx was confirmed by mass spectrometry based on the expected molecular weight.

### Western Blot Methylation Assays

Substrate mononucleosomes (800 nM) were incubated with S-adenosylmethionine (SAM; 250 µM), and reactions were initiated by the addition of wild-type or mutant G9a–GLP (1 µM). Reactions were carried out at 25 °C in 100 mM Tris-HCl (pH 8.0), 100 mM KCl, 1 mM MgCl₂, 20 µM ZnSO₄, and 10 mM β-mercaptoethanol (reaction buffer), and were quenched by the addition of Laemmli sample buffer.

Proteins were resolved by 18% SDS–PAGE and transferred to PVDF membranes using Tris–glycine transfer buffer containing 20% methanol. Membranes were blocked with INTERCEPT PBS blocking buffer (LiCor) and probed with antibodies against H3K9me2 (Abcam, ab1220) or H3K9me1 (Sigma-Aldrich, AB_2793303). An anti-H4 antibody (Active Motif, AB_2636967) was used as a loading control.

### G9a–GLP Histone Methyltransferase Assays

Methyltransferase activity of wild-type or mutant G9a–GLP was quantified by monitoring S-adenosyl-L-homocysteine (SAH) production using the MTase-Glo™ Methyltransferase Assay Kit (Promega), according to the manufacturer’s instructions. Reactions were performed at 25 °C in reaction buffer, supplemented with S-adenosylmethionine (SAM; 250 µM).

For kinetic analyses, concentrations of H3K9-containing substrates were varied while enzyme and SAM concentrations were held constant. Histone peptide substrates (0–500 µM) were assayed using 100 nM G9a–GLP, whereas mononucleosome substrates (0–3 µM) were assayed using 500 nM G9a–GLP. Peptide methylation reactions were incubated for 1 min and terminated by addition of 0.5% trifluoroacetic acid (TFA). Nucleosome methylation reactions were quenched after 30 min by the addition of 0.5% TFA to ensure measurements within the linear range of the reaction.

All methylation reactions were performed under multiple-turnover conditions. Initial velocities were determined from the linear portion of time-course measurements and plotted as a function of substrate concentration. Data obtained from duplicate or triplicate measurements were fit to the Michaelis–Menten equation, 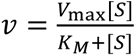 using GraphPad Prism to extract *K_M_*, *k*_cat_, catalytic efficiency (*k*_cat_/*K_M_*), and 95% confidence intervals. For direct determination of specificity constants (*k*_cat_/*K*_*M*_; Fig. 1), reactions were conducted at substrate concentrations below *K*_*M*_, where the slope of *v* versus [*S*] approximates 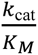. Luminescence was measured using a BioTek Cytation *K*_*M*_ 5 microplate reader, and SAH concentrations were calculated from a standard curve generated in parallel.

### Electrophoretic Mobility Shift Assays

Core nucleosomes (15 nM) were incubated with G9a–GLP or G9a–GLP mutants (0–5 µM) and S-adenosylmethionine (SAM, 250 µM) at 25 °C for 30 min in buffer containing 20 mM HEPES-NaOH (pH 7.5), 50 mM NaCl, 1 mM dithiothreitol, 0.03% NP-40, and 0.5% glycerol. Glutaraldehyde was then added to a final concentration of 0.1%, and the mixture was incubated at 4 °C for 30 min. The crosslinking reaction was quenched by adding Tris-HCl (pH 7.5) to a final concentration of 40 mM, followed by incubation on ice for 10 min. Samples were analyzed using 5% non-denaturing polyacrylamide gel electrophoresis in 0.5× TBE buffer and visualized with SYBR Green staining. K_1/2_ values (1-Fraction unbound) were calculated as before (55) by quantifying gel band intensities using Image Lab software (v5.2.1; Bio-Rad) and analyzed using GraphPad Prism.

### Crosslinking Mass Spectrometry

G9a–GLP (5 µM) was mixed with core nucleosomes (500 nM) and S-adenosylmethionine (SAM, 250 µM) in reaction buffer containing 20 mM HEPES-NaOH (pH 7.5), 50 mM NaCl, 1 mM dithiothreitol, 0.03% NP-40, and 0.5% glycerol, and incubated at 25 °C for 30 min. The sample was then crosslinked with disuccinimidyl sulfoxide (DSSO) at a final concentration of 800 µM and incubated at 4 °C for 30 min. The crosslinking reaction was quenched by the addition of Tris-HCl (pH 7.5) to a final concentration of 40 mM, followed by incubation at 25 °C for 10 min. Proteins were precipitated by acetone, and the pellet was alkylated with iodoacetamide and digested with trypsin. Peptides were desalted on a 100 μl Omix C18 tip (Agilent), dried, and reconstituted in 100 μl of 0.1% formic acid. Crosslinked peptides were fractionated by size-exclusion chromatography (SEC) on a Superdex Peptide column (56). Six SEC fractions were collected. The eluate was dried and reconstituted in 7.5 µl 0.1% formic acid. One third of each sample was used for mass spectrometry analysis.

Mass spectrometry was performed on an Orbitrap Exploris 480 equipped with an EasySpray nanoESI source, an EasySpray 75 μm × 15 cm C18 column coupled with an UltiMate 3000 RSLCnano system (Thermo Scientific). Each sample was loaded at 2% B at 600 nl/min for 35 min followed by a multisegment elution gradient to 35% B at 200 nl/min over 70 min with the remaining time used for column washing and reequilibration (buffer A: 0.1% formic acid (aq); buffer B: 0.1% formic acid in acetonitrile). Precursor ions were acquired at 120,000 resolving power, and ions with charges 3 to 8+ were isolated in the quadrupole using a 1.6 m/z unit window and dissociated by stepped HCD collision energy at 22, 30, 38% NCE. Product ions were measured at 30,000 resolving power.

Peak lists were generated using PAVA (in house Python app), searched with Protein Prospector v6.5.0 (57), and classified as unique residue pairs using Touchstone (an in-house R library) at 1% FDR and then further summarized and presented as domain-domain pairs using Touchstone. A custom database consisting of the human G9a-GLP constructs and histones was used for searching. Lysine analogs at H3K9 were encoded using unnatural amino acids to represent the Ecx, Nle, and Me2K residues in the software. A decoy database that was 10× longer than the target database was used in the Prospector search. Other search parameters included using tryptic specificity with 2-missed cleavages and tolerance of 10/20 ppm (precursor/product). DSSO* cross-linking was specified along with DSSO* dead-end modification at internal lysine residue and protein N-termini, methionine oxidation, N-terminal glutamate conversion to pyroglutamate, and incorrect monoisotopic precursor annotation were used as variable modifications. Three modifications per peptide were allowed and carbamidomethylation of cysteine was used as a contstant modification.

### Peptide MALDI-TOF analysis

G9a-GLP and mutants were prepared as above. Reactions were performed in reaction buffer with 250µM SAM and 100µM of and H3 tail peptide (amino acid 1-15; H3^1-15^, predicted mass by MS-isotope, UCSF prospector, 1560.9d) and 5µM enzyme. Reactions were carried out 25 °C following pre-equilibration. Reactions were started by addition of peptide and time courses stopped by addition of excess S-adenosyl homocysteine (SAH) to 1mM. For experiment 1, time points were 0’, 15’, 45’ and for experiment 2 0’ 5’ and 10’. Reactions were desalted using C18 ZipTips (MilliPore, #ZTC18S096). Data were collected on a Shimadzu Axima Performance. H3K9me0, me1, and me2 masses were confirmed via peptide H3^1-15^ (K9me0, me1, or me2) standards. External calibration was used with peptide standards. Data was acquired in linear mode.

### Structural modeling of the G9a–GLP heterodimer

The structural model of the G9a–GLP heterodimer was generated using AlphaLink2 (36), an AlphaFold2-based integrative modeling approach that incorporates cross-linking mass spectrometry (XL-MS)–derived distance restraints. All experimentally identified, high-confidence (SVM score > 2.34) heterotypic G9a–GLP cross-links were used as input restraints to guide heterodimer assembly and define the interaction interface. The resulting model was evaluated based on its consistency with the XL-MS restraints and overall structural plausibility. Cross-links were then mapped onto the model to validate the agreement between the predicted structure and the experimental data.

### Cryo-EM Sample Preparation

Asymmetric nucleosomes containing H3K9Nle/H3K9me2 modifications on 185 bp DNA were used. Nucleosomes (900 nM) were incubated with G9a–GLP (4 µM) and S-adenosylmethionine (SAM; 250 µM) in 20 mM HEPES-NaOH (pH 7.5), 40 mM KCl, 1 mM MgCl₂, 1 mM DTT, 0.03% NP-40, and 0.5% glycerol for 30 min at 25 °C. Crosslinking was initiated by adding glutaraldehyde to 0.1% and incubating at 4 °C for 30 min, followed by quenching with 50 mM Tris-HCl (pH 7.5) on ice for 10 min. Samples were purified by size-exclusion chromatography on a Superdex 200 Increase 10/300 column equilibrated in 10 mM Tris-Cl (pH 7.8), 50 mM KCl, and 1 mM DTT. Complex formation was verified by 5% non-denaturing polyacrylamide gel electrophoresis in 0.5× TBE and visualized with SYBR Green staining.

### Cryo-EM grid preparation and data collection

Nucleosome complex sample was kept on ice after concentration and directly loaded onto the glow-discharged holey carbon grid (Quantifoil 300 mesh Cu R1.2/1.3). The humidity in the chamber was kept at 95% and the temperature at 10°C. After 3 seconds of blotting, grids were plunge-frozen in liquid ethane using Thermo Scientific Mark IV vitrobot system. Electron micrographs were recorded on 300k eV Titan Krios electron microscope with a K3 electron detector at the Cryo-EM facility at St. Jude Childrens’s Research Hospital. Image pixel size was 0.6485 Å per pixel on the object scale. Data were collected in a defocus range of 9,000–30,000 Å with a total exposure of 80 e-Å. In total, 170,000 frames were collected. Frames were aligned with the MotionCor2 software (58). The contrast transfer function parameters were determined using CTFFIND4 (59). Particles were picked by Topaz (60) using pre-trained model. Particles were windowed and 2D class averages were generated with the Relion software package (61). Inconsistent class averages were removed from further data analysis. The initial reference was filtered to 40 Å in Relion. C1 symmetry was applied during refinements for all classes. Particles were split into 2 datasets and refined independently, and the resolution was determined using the 0.143 cut-off (Relion auto refine option). All maps were filtered to resolution using Relion with a B-factor determined by Relion.

### Model building and refinement

Molecular models were built using ChimeraX (62) and Coot (63). To build the model of nucleosome complex with G9a Ankryin domain, the pdb model of Widow 601 nucleosome ((35) PDB: 1AOI) and Alphafold3 (64) predicted G9a Ankryin was built by rigid-body fitting and refined into the cryo-EM map. To build the model of nucleosome complex with G9a-GLP dimer, the XL-MS model of G9a-GLP dimer was fit into the cryo-EM structure in ChimeraX.

## Supporting information

All Supplemental Figures and Legends.

## Acknowledgements

We thank Dr. Jo Hyunil for support with MALDI-TOF experiments, and members of the Al-Sady and Buchwalter labs for critical feedback. B.A.-S. was supported by NIH award R35GM141888 and a Melanoma Research Alliance Pilot Award 1264883. M.H. was supported by St. Jude Children’s Research Hospital, the American Lebanese Syrian Associated Charities and NIH awards R01GM135599 and R35GM158165. B.A.S. and M.H. were supported by 5R21DA06282. Y.D. was supported by a Gephardt Postdoctoral Fellowship from the Cancer Biology Program. P.A.C. was supported by NIH grant GM149229. B.P. was supported by NIH grant R01GM128431. B.P. and B.A.S. were supported by the Sandler Program for Breakthrough Biomedical Research. S.D.W. was supported by fellowships from American Cancer Society (PF-20-105-01-DMC) and the Charles King Trust. E.S. was supported by a fellowship from the Ford Foundation. Mass spectrometry experiments were supported by the Adelson Medical Research Foundation and the University of California, San Francisco Program for Breakthrough Biomedical Research.

