## Supplementary material for "Histone 3 lysine 9 dimethylation by the G9a-GLP heterodimer requires intranucleosomal product reading": All Supplemental Figures and Legends.

#### Supporting material.

**Supporting Figure 1: G9a-GLP exhibits only minor kinetics differences on H3K9me0 or H3K9me1 peptides. A.** Michaelis-Menten curve for G9a-GLP and H3 tail peptides (H3$^{1-15}$K9me0 and H3$^{1-15}$K9me1). Error bars indicate standard deviation of 3 repeats. **B.** Kinetic constants for methylation of H3$^{1-15}$K9me0 and H3$^{1-15}$K9me1 by G9a-GLP.

**Supporting Figure 2: G9a-GLP mutants mono- and dimethylate H3 peptides.** Methylation and MALDI-TOF Experiment 1: comparison of wildtype G9a-GLP (A.), ∆ANK (B.) and G9a-GLP$^{SWAP}$ (C.). The reaction was run for 15' and 45'. The 0', 15' and 45' traces are shown overlapped. 0', black; 15', orange; 45' , green. 45' times points show predominately H3$^{1-15}$K9me2. Methylation and MALDI-TOF Experiment 2: comparison of wildtype G9a-GLP (D.), G9aANK$^{m}$-GLPANK$^{m}$ (E.) and G9aANK8$^{GS}$-GLP (F.). The reaction was run for 5' and 10'. The 0', 5' and 10' traces are shown overlapped. 0', black; 5', blue; 10' , yellow. The measured mass for the H3$^{1-15}$K9me0 at 0' in each run is shown peptide. One of two repeat spectra is shown per timepoint.

**Supporting Figure 3. crosslink mass spectrometry with initial substrate and reaction intermediates. A.** Overview of catalytic histone lysine mimics ethylcysteine (EcX), Norleucine (Nle) and methionine (Met) **B.** Nondenaturing gel electrophoresis of nucleosome complexes formed by H3K9Ecx (Ecx), H3K9Nle (Nle), and H3K9Met (Met) nucleosomes with G9a–GLP. Binding reactions were performed with nucleosomes (20 nM) and G9a–GLP (5.6 µM) **C.** Overview of all DSSO crosslinks between and within reaction intermediate nucleosome and G9a-GLP. **D.** As in C. but for initial substrate nucleosome.

**Supporting Figure 4: G9aANK8$^{GS}$-GLP binds poorly to nucleosomes. A.** sequence of the ANK8 region on G9a and GLP. Unique region boxed. **B.** EMSA with substrate mimic (black dots) and H3K9me1 (teal dots) nucleosomes (15nM) and G9aANK8$^{GS}$-GLP. G9aANK8$^{GS}$–GLP was titrated in a seven-point serial dilution from 6.25 μM to 0 μM (6.25, 3.13, 1.56, 0.78, 0.39, 0.20, and 0 μM).

**Supporting Figure 5. Complex preparation for cryo-EM, class averages, and resolution of electron density maps. A.** Production and isolation of the G9a-GLP:reaction intermediate (H3K9Nle/H3K9me2) nucleosome for cryo-electron microscopy. Crosslinked nucleosome : G9a-GLP complexes were purified by FPLC. Single or pooled indicated fractions were concentrated; the final material is shown on the right. Lanes indicated by a green dot were spotted on grids. **B.** Representative cryo-EM micrograph and 2D classes of H3K9Nle/H3K9me2 nucleosome complex with G9a-GLP. **C.** TOP: Top view of reaction intermediate nucleosome with G9a-GLP, electron density map 709. BOTTOM: Fourier shell correlation (FSC) curve showing the resolution of map 709. **D.** TOP: Top view of reaction intermediate nucleosome with G9a-GLP, electron density map 808. BOTTOM: Fourier shell correlation (FSC) curve showing the resolution of map 808.

**Supporting Figure 6: CLMS of H3 tail to nucleosome. A.** Diagram for intranucleosomal crosslinks anchored on the H3 tail for substrate mimic (H3K9EcX/H3K9EcX) nucleosome from the CLMS experiment of the G9a-GLP:nucleosome complex. **B.-D.** crosslinks between the H3 tail and H2A, H2B, and H4 histones in the complex. **E.** Diagram for intranucleosomal crosslinks anchored on the H3 tail for reaction intermediate mimic (H3K9Nle/H3K9me2) nucleosome from the CLMS experiment of the G9a-GLP:nucleosome complex. **F.-G.** As in B.-D. but with reaction intermediate mimic (H3K9Nle/H3K9me2) nucleosome in the presence of G9a-GLP. Substrate mimic-specific crosslinks, skyblue; reaction intermediate-specific crosslinks, red.

**Supporting Figure 7: AlphaLink2 model of G9a-GLP. A.** Crosslinks inside the G9-GLP dimer derived from CLMS experiments in Figure 3 (with substrate and reaction intermediate mimic nucleosomes). Shown are the Domain-level bubble plots of G9a–GLP interactions. **A.** G9a-GLP intramolecular crosslinks in the substrate mimic experiment. **B.** As in A., but for a reaction intermediate mimic experiment. Unique crosslinks in B. highlighted with red circles. The sizes of circles reflect the strength of interaction (CSM sum), and the color indicates the prevalence of interaction. **C.** AlphaLink2 structural model of the G9a–GLP heterodimer colored by predicted Local Distance Difference Test (pLDDT; blue, high confidence; red, low confidence). **D.** pLDDT plot of C. **E.** Predicted aligned error (PAE) matrix for the AlphaLink2 model, indicating high confidence in the relative positioning of the interacting G9a and GLP domains. Crosslinks satisfied (red) and not satisfied (blue) in the model are indicated. X-axis numbers indicate amino acids of the combined G9a-GLP ANK-SET dimer, from arranged form the N-terminal residue of G9a ANK to the C-terminal residue of GLP SET. **F.** Overlay of AlphaLInk2 and AlphaFold3 (AF3) models.

#### SFigure 1

**A**

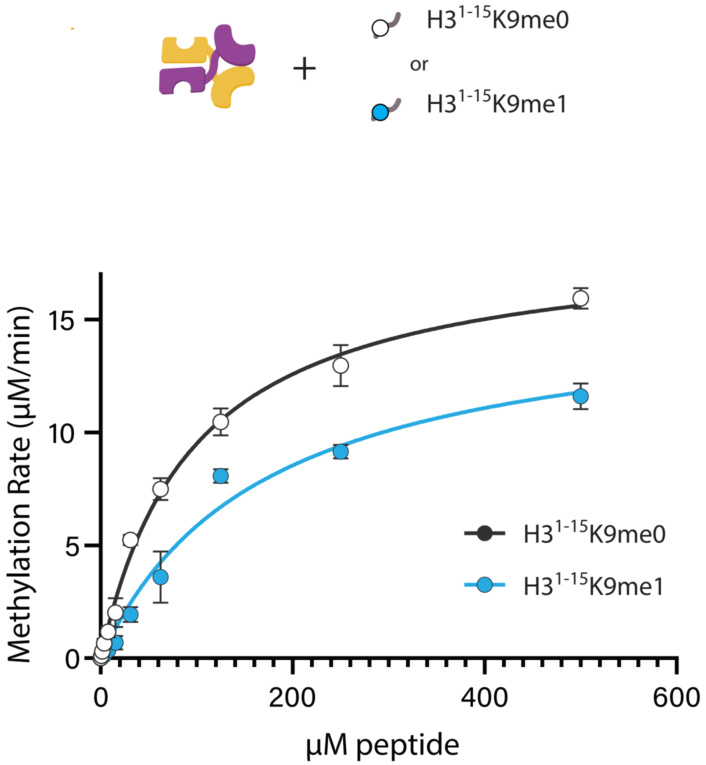


**B**

| G9a-GLP | $K_M$ (μM) | $k_{cat}$ (min$^{-1}$) | $\frac{k_{cat}}{K_M}$ (μM$^{-1}$ min$^{-1}$) |
|---|---|---|---|
| H3$^{1-15}$K9me0 | 96.2 (95% CI 84.6-110) | 186 (95% CI 177.6-195.6) | **1.93** |
| H3$^{1-15}$K9me1 | 169.6 (95% CI 132.4-219.6) | 157.2 (95% CI 142.2-176.4) | **0.92** |

**SFigure 2**

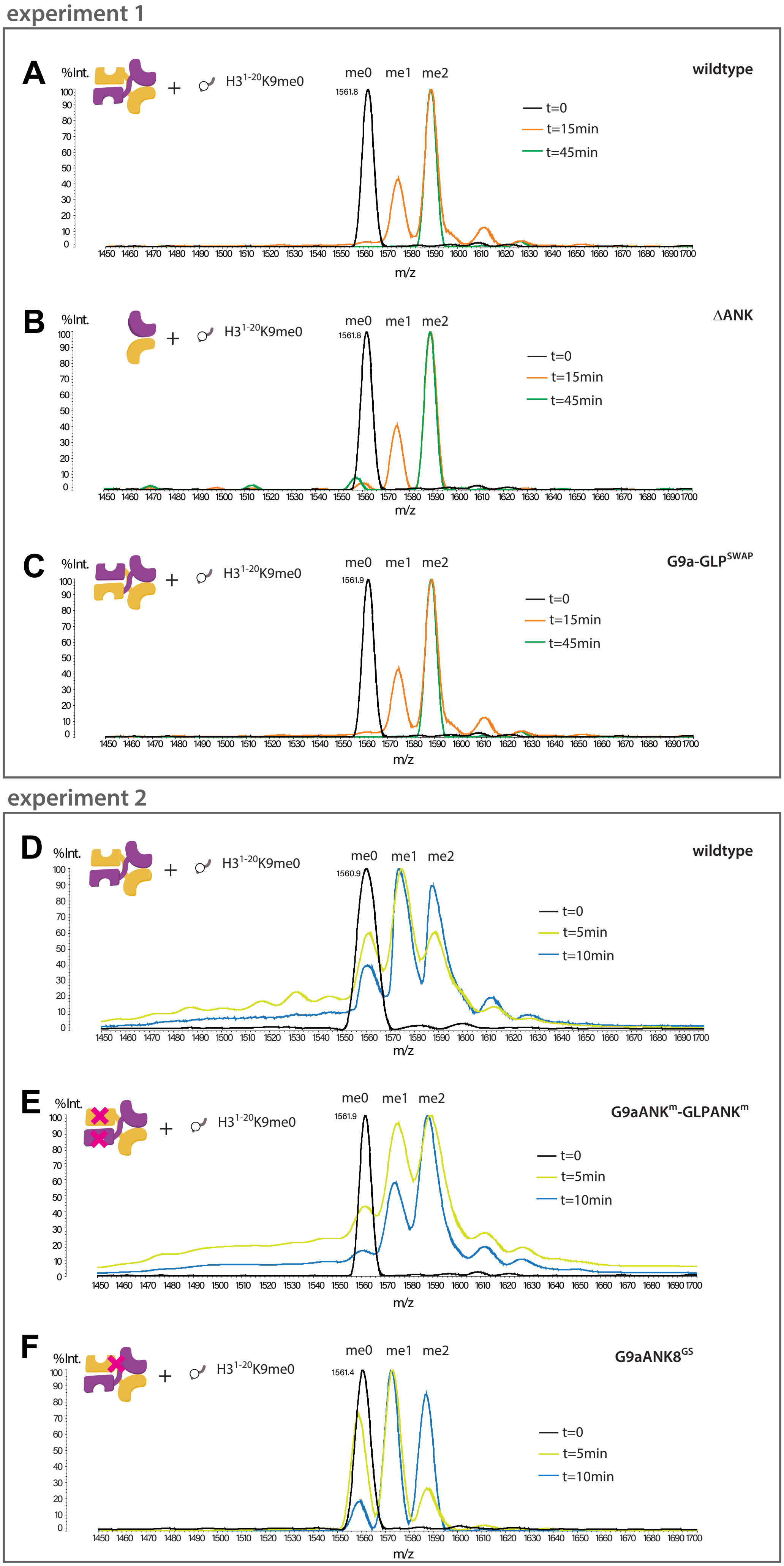

experiment 1
A
%Int.
H3 1-20 K9me0
me0
me1
me2
1561.8
wildtype
t=0
t=15min
t=45min
m/z
B
ΔANK
C
1561.9
G9a-GLP SWAP
experiment 2
D
1560.9
t=5min
t=10min
E
G9aANK m -GLPANK m
F
1561.4
G9aANK8 GS

### SFigure 3

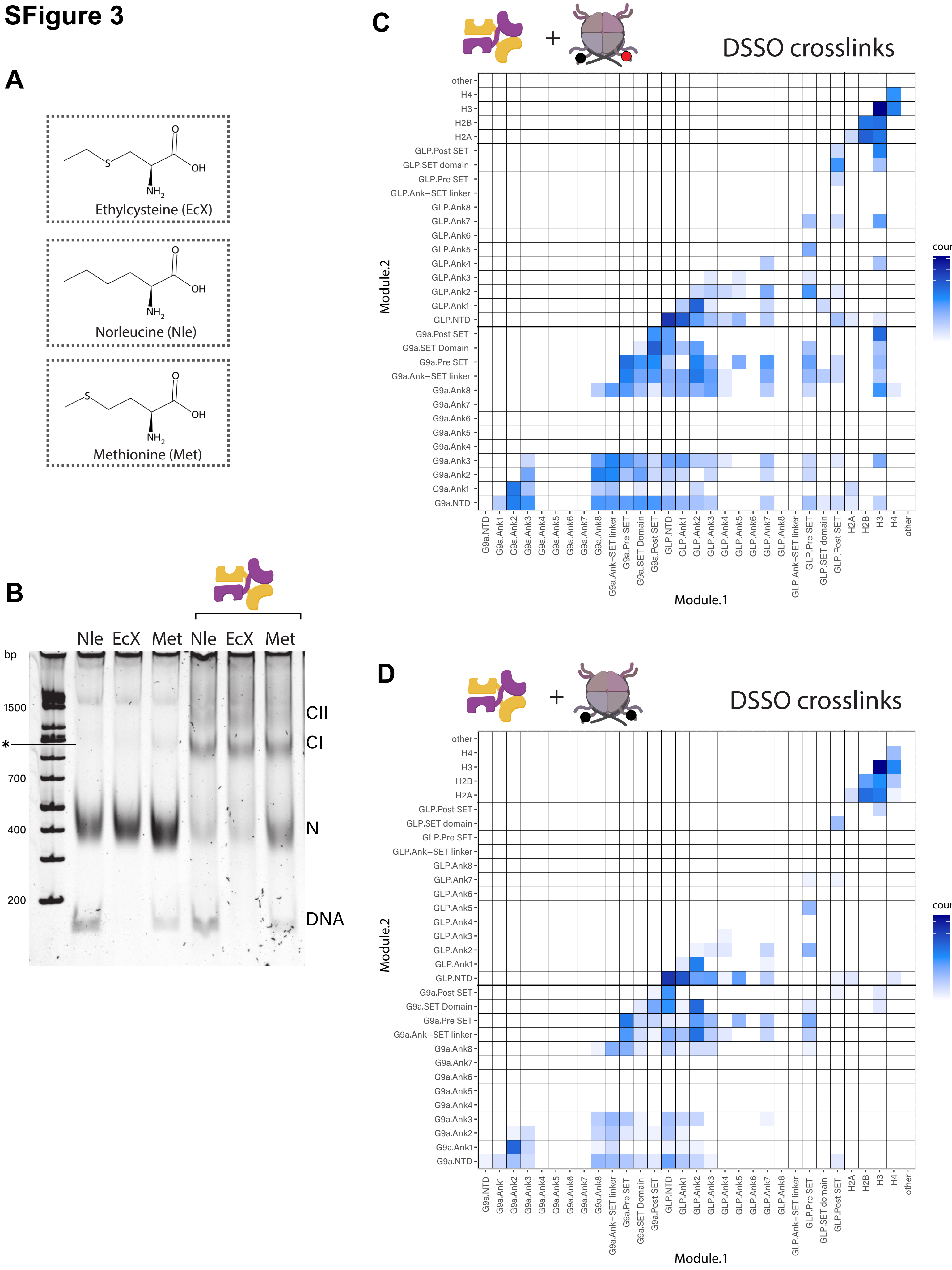
A
Ethylcysteine (EcX)
Norleucine (Nle)
Methionine (Met)
O
OH
S
NH2
B
Nle EcX Met Nle EcX Met
bp
1500
*
700
400
200
CII
CI
N
DNA
C
DSSO crosslinks
+
Module.2
Module.1
counts
400
300
200
100
0
other
H4
H3
H2B
H2A
GLP.Post SET
GLP.SET domain
GLP.Pre SET
GLP.Ank−SET linker
GLP.Ank8
GLP.Ank7
GLP.Ank6
GLP.Ank5
GLP.Ank4
GLP.Ank3
GLP.Ank2
GLP.Ank1
GLP.NTD
G9a.Post SET
G9a.SET Domain
G9a.Pre SET
G9a.Ank−SET linker
G9a.Ank8
G9a.Ank7
G9a.Ank6
G9a.Ank5
G9a.Ank4
G9a.Ank3
G9a.Ank2
G9a.Ank1
G9a.NTD
D
DSSO crosslinks
+
Module.2
Module.1
counts
200
100
0

#### SFigure 4

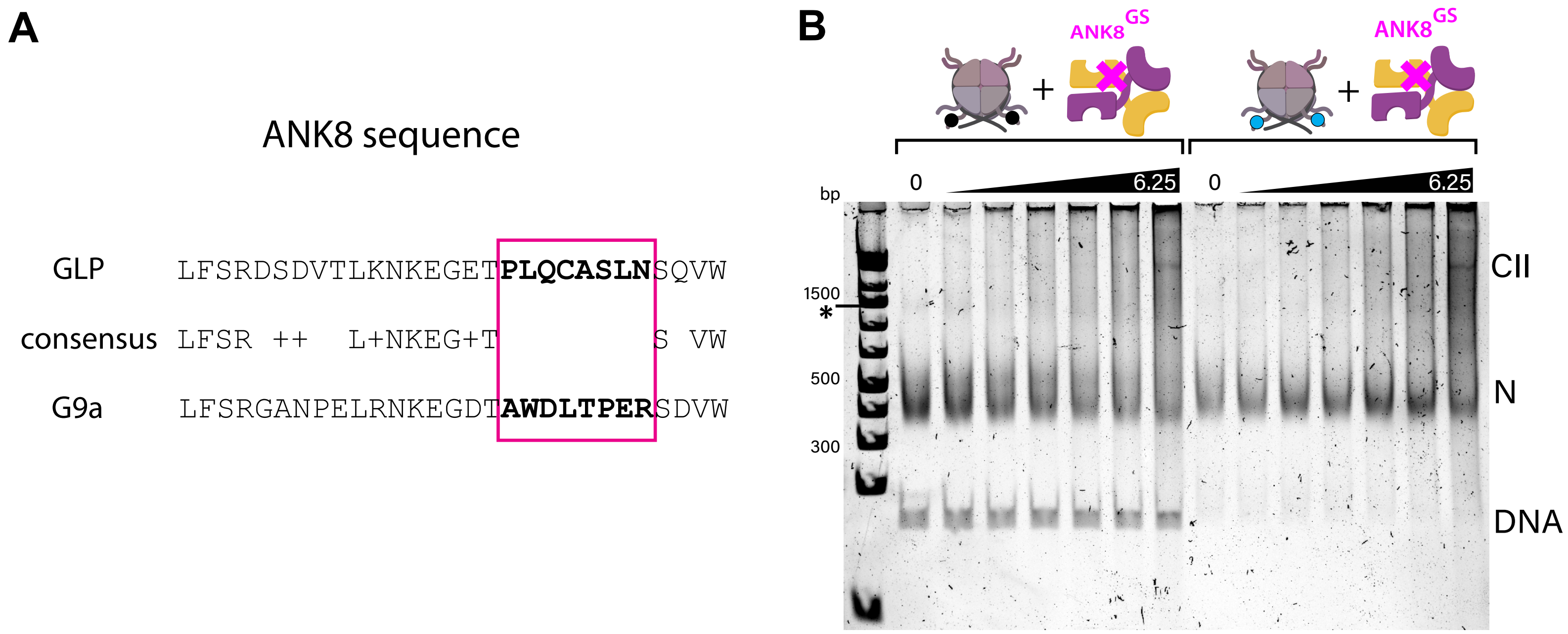
A
ANK8 sequence
GLP LFSRDSDVTLKNKEGETPLQCASLNSQVW
consensus LFSR ++ L+NKEG+T S VW
G9a LFSRGANPELRNKEGDTAWDLTPERSDVW
B
ANK8GS
ANK8GS
0
6.25
0
6.25
bp
1500
*
500
300
CII
N
DNA

### SFigure 5

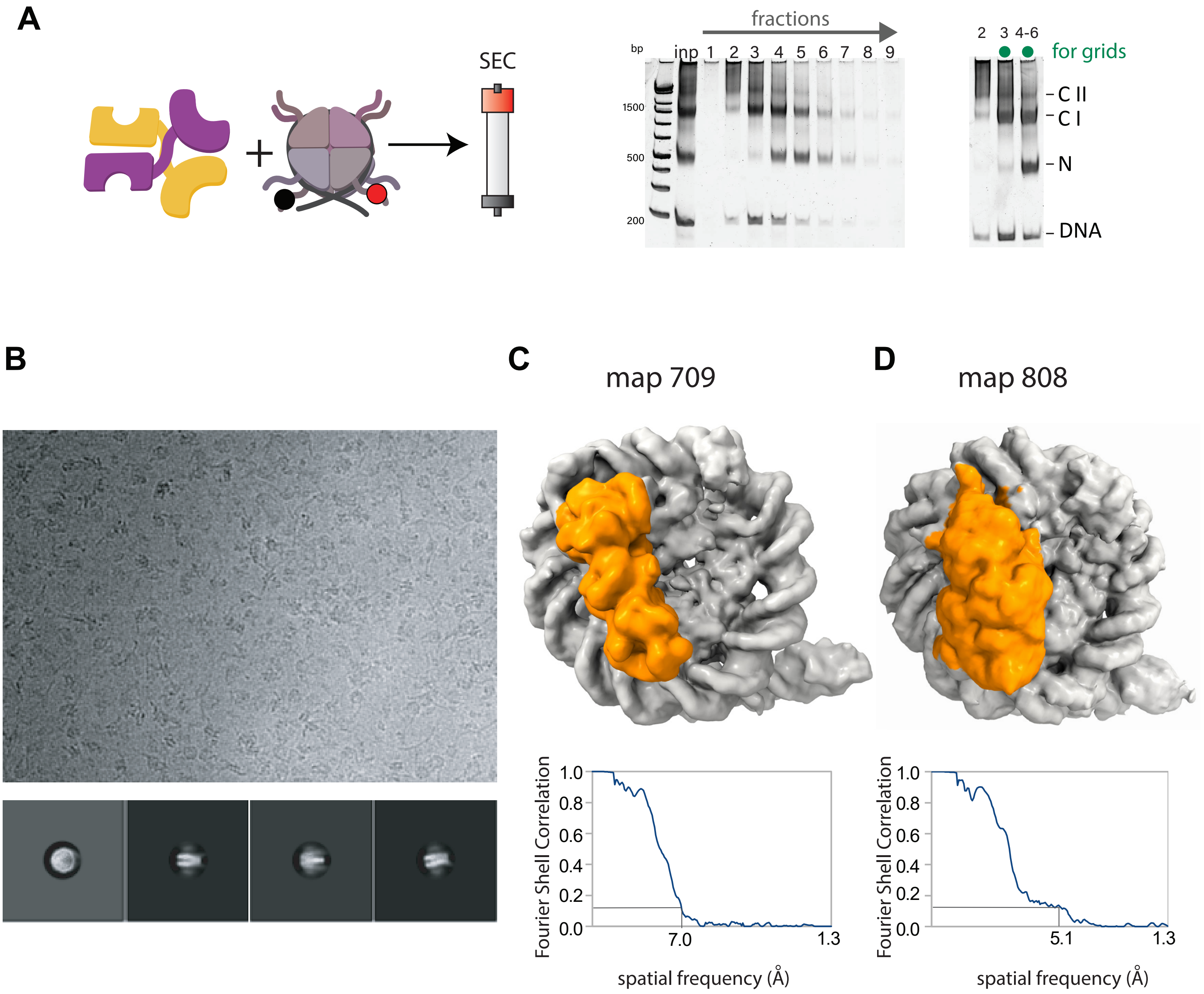
A
SEC
fractions
bp
inp
1 2 3 4 5 6 7 8 9
1500
500
200
2 3 4-6
for grids
C II
C I
N
DNA
B
C
map 709
Fourier Shell Correlation
1.0
0.8
0.6
0.4
0.2
0.0
7.0
1.3
spatial frequency (Å)
D
map 808
Fourier Shell Correlation
1.0
0.8
0.6
0.4
0.2
0.0
5.1
1.3
spatial frequency (Å)

### SFigure 6

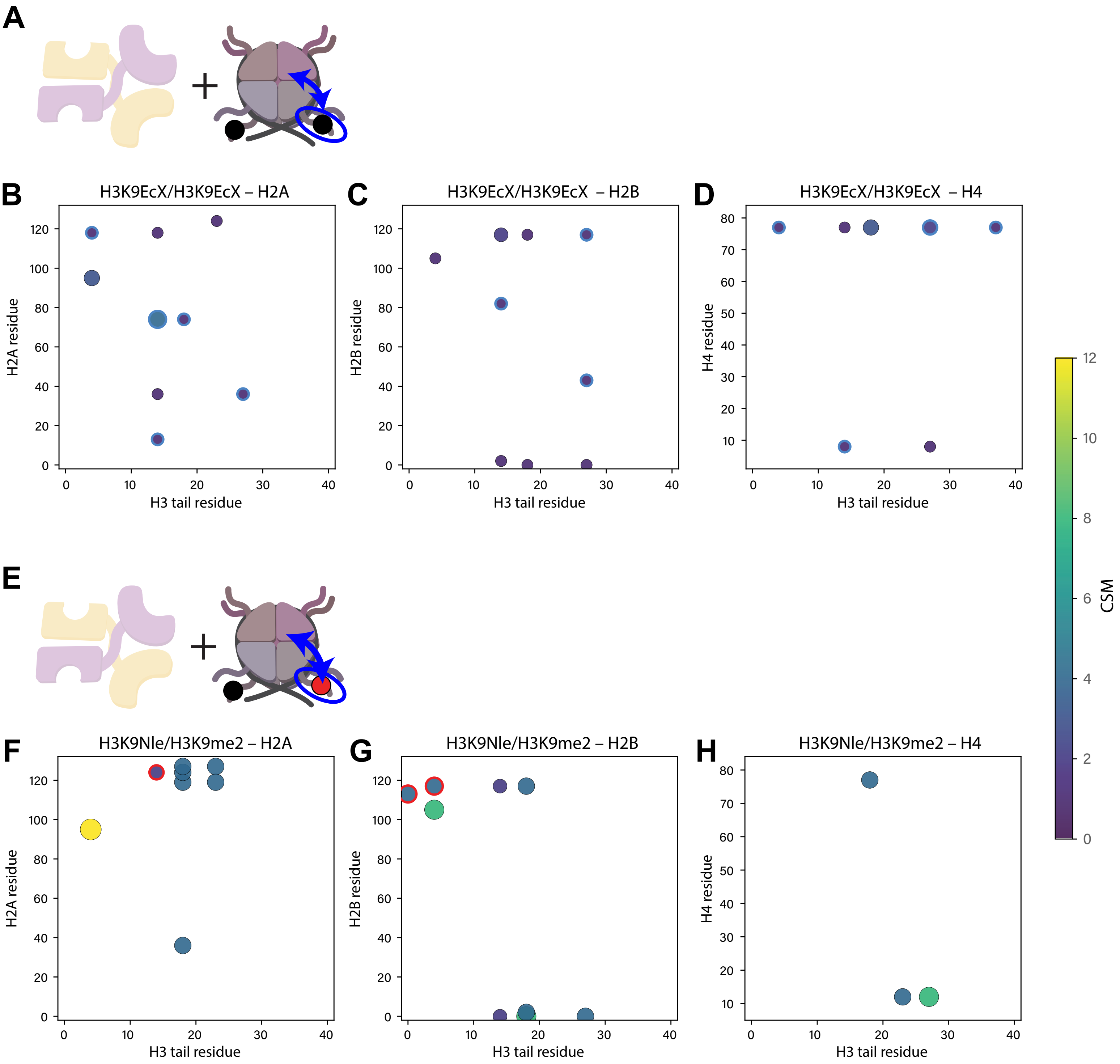

A
B
H3K9EcX/H3K9EcX – H2A
H2A residue
H3 tail residue
C
H3K9EcX/H3K9EcX – H2B
H2B residue
H3 tail residue
D
H3K9EcX/H3K9EcX – H4
H4 residue
H3 tail residue
E
F
H3K9Nle/H3K9me2 – H2A
H2A residue
H3 tail residue
G
H3K9Nle/H3K9me2 – H2B
H2B residue
H3 tail residue
H
H3K9Nle/H3K9me2 – H4
H4 residue
H3 tail residue
CSM

### SFigure 7

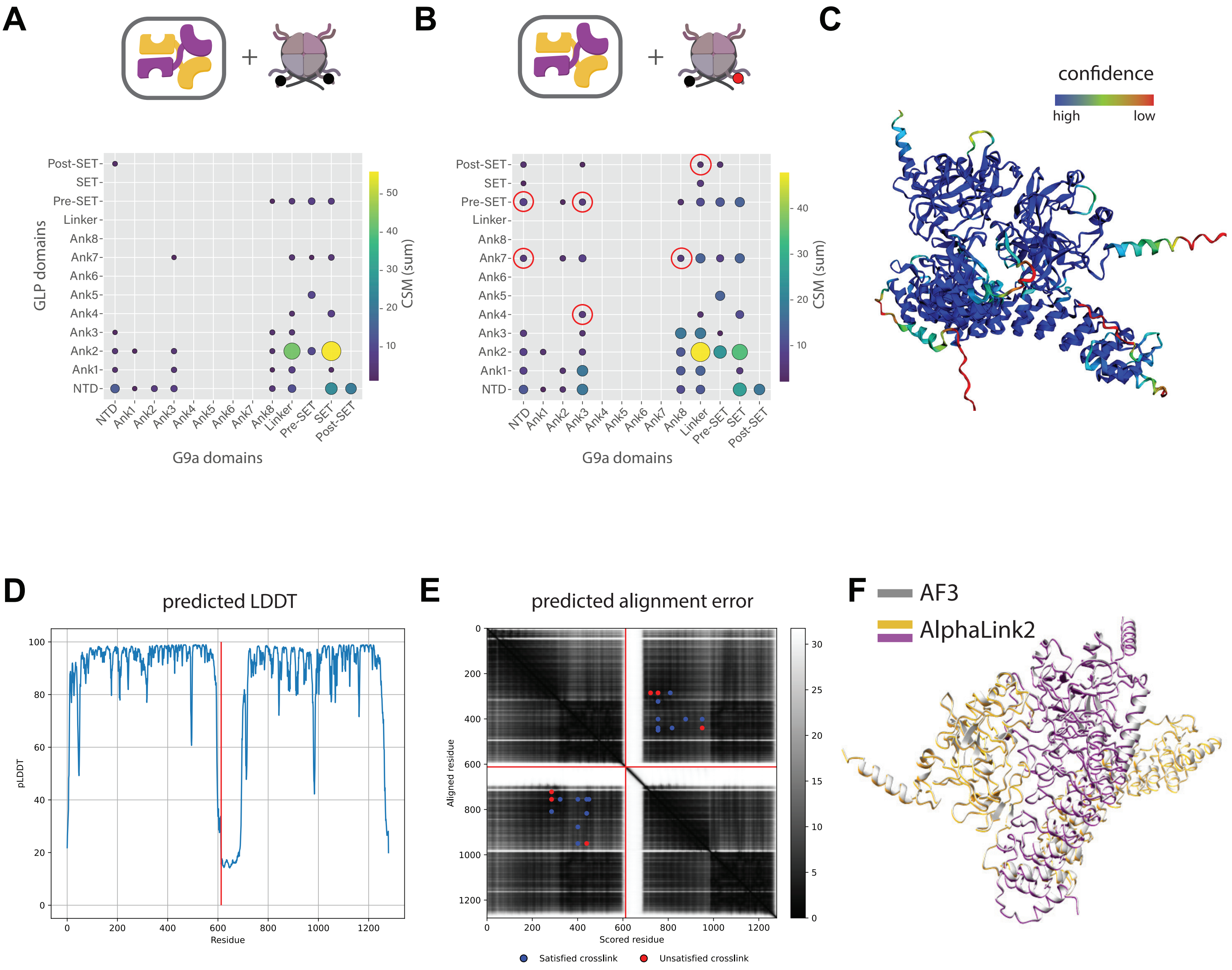
A
GLP domains
Post-SET
SET
Pre-SET
Linker
Ank8
Ank7
Ank6
Ank5
Ank4
Ank3
Ank2
Ank1
NTD
G9a domains
CSM (sum)
B
C
confidence
high
low
D
predicted LDDT
pLDDT
Residue
E
predicted alignment error
Aligned residue
Scored residue
Satisfied crosslink
Unsatisfied crosslink
F
AF3
AlphaLink2
